# Single-cell metabolomics reveals infection-specific metabolic reprogramming of human macrophages

**DOI:** 10.64898/2026.09.03.749052

**Authors:** Paula Martinez-Oca, Arnaud Meng, Quentin Vanbellingen, Marvin Albert, Antoine Thomas, Perrine Bomme, Francisco-Javier Garcia-Rodriguez, COSIPOP Study Group, Jean-Yves Tinevez, Sandrine Aros, Carmen Buchrieser, Pedro Escoll

## Abstract

During intracellular infection, host cells adopt different metabolic states that traditional bulk analyses cannot distinguish. Using single-cell spatial metabolomics of human macrophages infected with *Legionella pneumophila*, we show that bacterial uptake activates host metabolism, whereas the bacterial effectors secreted through the type IV secretion system counteract this response and promote a glycolytic shift. The activity of effectors also generates distinct metabolic states within the infected macrophage population.

---

*Legionella pneumophila* injects over 300 bacterial effectors through its type IV secretion system (T4SS) to build a replicative vacuole in macrophages, extensively remodeling host metabolism ^1,2^. Whether individual macrophages respond uniformly or adopt heterogeneous metabolic states has remained inaccessible, as bulk metabolomics averages over millions of cells, and single-cell RNA sequencing cannot measure metabolites. We developed SpatialMetProfiler, a MALDI mass-spectrometry-imaging (MSI) pipeline that co-registers each single-cell metabolic spectrum with the corresponding infection status, as determined by bacterial mCherry fluorescence, in primary human monocyte-derived macrophages (hMDMs) (Fig. 1a, Extended Data Fig. 1). Thus, only infected cells are analyzed, eliminating the noise from uninfected bystanders that confounds bulk approaches. To separate the contributions of bacterial uptake, bacterial viability and T4SS effector delivery to the metabolic reprogramming of host cells, we profiled over 11,000 single cells across four infection conditions: non-infected (NI, baseline), heat-killed *L. pneumophila* (HK, uptake of non-viable bacteria and sensing of pathogen-associated molecular patterns (PAMPs), Extended Data Fig. 2), the T4SS-deficient *L. pneumophila* Δ*dotA* mutant (live, avirulent mutant lacking a functional T4SS necessary for effector translocation), and *L. pneumophila* wild-type strain Paris (WT, infection competent, clinical strain). We analyzed 5 h post-infection to capture the pre-replicative phase of infection, when T4SS effector translocation has started but bacterial replication has not yet commenced ^3,4^, providing a defined biological window in which host metabolic reprogramming can be attributed to effector activity rather than to differential bacterial replication between wells and strains. Across all four conditions, single-cell metabolic profiles formed a continuous, overlapping landscape without clear infection condition-specific clusters (Fig. 1b). However, two main clusters seem to emerge, where one mainly contains the metabolic profiles of NI cells and the other one mainly those of bacteria-infected cells. Interestingly, metabolic profiles of WT-infected cells can be found in both clusters and between the clusters indicating that infection shifts metabolism within a shared space rather than creating distinct populations. Comparing the infection conditions pairwise isolates three contributions to host metabolic reprogramming: T4SS effector delivery (WT vs Δ*dotA*), sensing of bacterial components (HK vs NI), and bacterial viability (live-but-avirulent versus dead bacteria, Δ*dotA* vs HK).

**Fig. 1.**
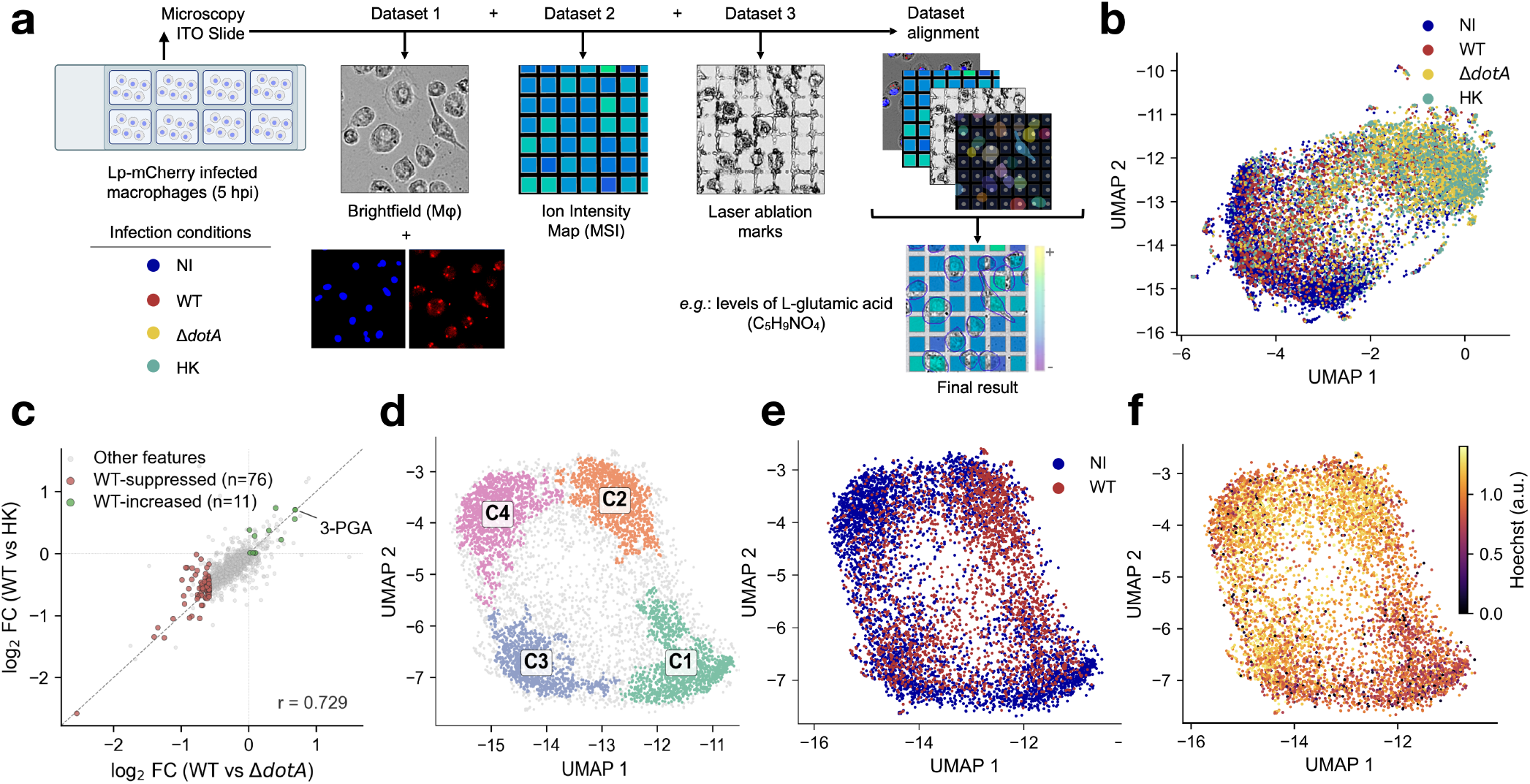
Single-cell spatial metabolomics separates the metabolic contributions of bacterial uptake, viability and T4SS effector delivery. (**a**) SpatialMetProfiler pipeline. Macrophages infected on an ITO slide (5 hpi) are imaged (brightfield, Hoechst, mCherry), MALDI ion-intensity maps and post-MALDI ablation marks are acquired, and the three datasets are aligned to assign each metabolite spectrum to single, segmented cells (*e.g*. L-glutamic acid). Lp = *L. pneumophila*; M*φ* = macrophages; ITO = Indium Tin Oxide; MSI = Mass Spectrometry Imaging. (**b**) Global UMAP of all single-cell spectra across the four conditions. (**c**) Per-feature log_2_ fold-change, WT vs Δ*dotA* (x) versus WT vs HK (y). The 76 features concordantly decreased in WT compared to Δ*dotA* and HK are highlighted (Pearson r = 0.73, identity line dashed). 3-phosphoglycerate (3-PGA, upper right) is the principal WT-increased feature. (**d**) HDBSCAN of the WT-NI UMAP (four subclusters). (**e**) WT vs NI UMAP (831 features) colored by infection status. (**f**) Same as e colored by normalized Hoechst signal.

How do T4SS effectors impact metabolic reprogramming? As the WT and its isogenic Δ*dotA* mutant are both internalized alive, but differ in effector delivery, their comparison helps us to better understand the contribution of early secreted effectors. Of 64 differentially abundant features (limma, FDR < 0.10; 1,606 features detected, 653 annotated), 62 were suppressed in WT infection (Extended Data Fig. 3). This result indicates that effectors impact the metabolic remodeling associated with bacterial uptake (246 features with increased relative abundance in avirulent Δ*dotA*; Extended Data Fig. 3). To test whether this suppression depends on the choice of avirulent reference, we modelled all four conditions jointly (shared-fit limma; Methods). Among features significantly increased upon uptake of Δ*dotA* or HK bacteria relative to NI, 76 were concordantly lower in WT than in both Δ*dotA* and HK, and 11 were higher (Fig. 1c). Among the 11 features increased in WT infection in this shared-fit analysis, 3-phosphoglycerate (3-PGA) was identified, consistent with our previous report that infection induces a Warburg-like glycolytic phenotype ^2^ that might divert carbon flow in the host cell to later support bacterial replication. Unsupervised HDBSCAN clustering on the WT-NI UMAP identified four metabolic subclusters (Fig. 1d). These clusters were not separated by cell shape (cytoplasm eccentricity) and therefore do not map onto the classical M1/M2 polarization axis ^5^, although they differed in nuclear staining (Fig. 1ef and Extended Data Fig. 4). Indeed, clusters C2 and C3, enriched for infected cells, showed higher Hoechst intensity, potentially reflecting infection-associated nuclear condensation and higher cellular stress, whereas C1, enriched in NI cells, displayed the lowest signal. This heterogeneity within the infected population may reflect variation in effector delivery between cells, variations in the timing of metabolic reprogramming, or an intrinsic metabolic plasticity of primary human macrophages that extends beyond the M1/M2 paradigm. Effector delivery also reshaped heterogeneity within the infected population, as seen in within-condition embeddings. WT resolved into five metabolic subclusters versus two for every other condition (Fig. 2a-d), indicating that effector delivery generates additional metabolic states rather than merely shifting the population as a whole. Two of these subclusters carried distinct pathway signatures (Extended Data Fig. 5). WT-C3 subcluster was associated with biosynthetic pathways (pentose phosphate pathway, cofactor and aminoacid biosynthesis, nicotinate/nicotinamide metabolism) and WT-C5 with 2-oxocarboxylic acid metabolism, which might be consistent with transamination and mobilization of host amino acids due to bacterial auxotrophies. Together, these results suggest that, although WT infection broadly dampened the metabolic activation otherwise triggered by bacterial uptake, the glycolytic intermediate 3-PGA selectively accumulated. WT infection also split this response into coexisting substates rather than a single uniform program.

**Fig. 2.**
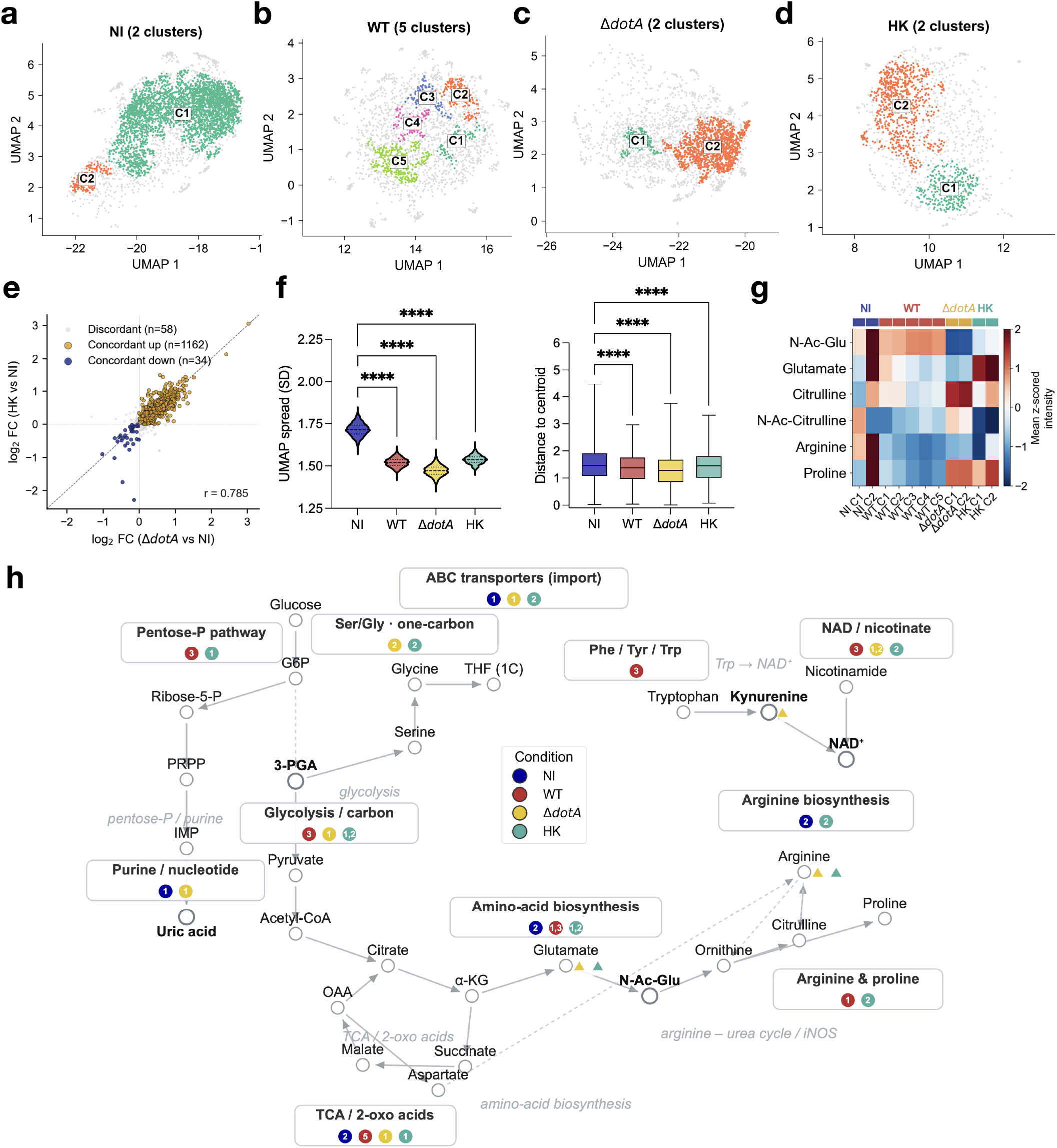
Infection converges macrophage metabolism while T4SS activity superimposes discrete substates. (**a-d**) Within-condition UMAP and HDBSCAN for NI, WT, Δ*dotA*, HK. (**e**) Per-feature log_2_ fold-change, Δ*dotA* vs NI (x) versus HK vs NI (y). 1,162 features concordantly up, 34 concordantly down, 58 discordant (Pearson r = 0.79). (**f**) Single-cell dispersion per condition (bootstrap UMAP spread, left; distance to population centroid, right). NI is the most dispersed, all bacterial conditions more constrained (Mann-Whitney U, P < 0.001), Δ*dotA* most homogeneous. (**g**) Arginine-pathway signature across within-condition subclusters (mean z-scored intensity of N-acetylglutamate, glutamate, citrulline, N-acetyl-citrulline, arginine and proline). (**h**) Integrated model of core macrophage metabolism summarizing all infection-resolved programs. Chips are color-coded according to infection conditions (legend in the center). Numbers inside chips are the corresponding clusters per infection condition depicted in Fig. 2a-d.

How does sensing bacterial components impact macrophage metabolism? Infection with heat-killed bacteria allows to preserve PAMPs and thus sensing of bacterial components without active bacterial metabolism and delivery of effectors. Compared to NI, HK bacteria elicited a broad host response (168 features increased, Extended Data Fig. 3) largely shared with the Δ*dotA* strain (Fig. 2e). 22 features were annotated, and pathway-level mapping was dominated by single features such as N2-succinyl-L-glutamic acid 5-semialdehyde (arginine and proline metabolism, higher in HK) or uric acid (purine metabolism), which was higher in NI (Extended Data Fig. 5). As *L. pneumophila* LPS signals through TLR2 rather than TLR4^6^, the HK response would be expected to resemble that of TLR2 agonists, which in human monocytes engage oxidative and lipid metabolism rather than the tricarboxylic-acid-cycle intermediates that accumulate upon TLR4 stimulation ^7^. Consistent with this, some of the annotated features increased in HK were lipid species (phosphatidylinositols), an acylcarnitine (fumarycarnitine), and an octadecatrienynoic acid, although the TCA intermediates characteristic of the TLR4 response were not detected in our dataset. The detected increase in phosphatidylinositol species PI(34:0) and PI(34:2) with uptake of HK bacteria, independently of bacterial effectors, is consistent with membrane remodeling during *L. pneumophila* phagocytosis ^8^. Bacterial-component sensing therefore drives broad activation of macrophage metabolism, in contrast to the attenuation associated with effectors.

Does the metabolic response to the uptake of live avirulent bacteria differ from the uptake of dead bacteria? Phagocytosing live but replication deficient bacteria, such as *L. pneumophila* Δ*dotA* broadly activated metabolism compared to NI (246 features increased, Extended Data Fig. 3), an activation similar to the HK response (Δ*dotA* vs NI and HK vs NI concordant across features, r = 0.79; Fig. 2e). The direct Δ*dotA*-versus-HK comparison (live versus dead, neither delivering effectors) revealed that only one feature was different (Extended Data Fig. 3), and Δ*dotA* formed the most homogeneous population of all conditions (Fig. 2c,f). Bacterial viability (Δ*dotA versus* HK) therefore contributes only little to host metabolic reprogramming consistent with uptake and PAMP sensing. However, T4SS effector activity induces a distinct attenuation of the metabolic response.

Taken together, these results of single-cell spatial metabolomics show that infection reshaped metabolic heterogeneity in two opposing ways. First, macrophages exposed to bacteria became more similar to one another than resting macrophages were, lying closer to the center of their own population whether the bacteria were virulent, avirulent or heat-killed (distance to population centroid, Mann–Whitney U, P < 0.001 for each, Fig. 2f). Second, and in the opposite direction, WT infection generated additional metabolic states. WT macrophages resolved into five subclusters, whereas every other condition resolved into two (Fig. 2a-d). This extra substructure most likely reflects stochastic cell-to-cell variation in effector delivery ^9,10^ or in how far reprogramming has progressed, rather than pre-existing differences between macrophages, which would otherwise appear equally in NI cells. Such variation may help explain why some macrophages restrict bacterial growth while others become permissive ^4^, as replication would require that the metabolic state of the host cell matches the nutritional demands of the bacterium. Arginine-pathway metabolites also distinguished subpopulations before and after infection. While resting macrophages contained a discrete arginine-biosynthesis subpopulation (NI-C2), after bacterial encounter citrulline (a product of the inducible nitric-oxide-synthase reaction) accumulated selectively in Δ*dotA* subclusters and glutamate in HK subclusters, while proline (generated by the alternative, arginase-dependent arm of arginine catabolism) was elevated in both relative to WT (Fig. 2g). As a pre-loaded arginine reservoir has been proposed to poise innate cells for activation and to support trained immunity ^11^, it is tempting to speculate that such pre-loaded resting state of NI subcluster could be the metabolic precursor from which infection-associated arginine substates emerge. Testing this hypothesis would require time-resolved metabolic tracking of individual living cells. Finally, we summarized the condition-resolved programs in an integrated map of core macrophage metabolism (Fig. 2h). This approach can now be used to test whether broad effector-mediated metabolic attenuation is a conserved virulence strategy across intracellular pathogens.

## Acknowledgements

We thank all members of the Buchrieser laboratory for fruitful discussions. We thank Nathalie Aulner, Nassim Mahtal, Julien Fernandes and the Photonic BioImaging (PBI) UTechS at Institut Pasteur for their support. We are grateful to the healthy volunteers for their participation in the study, and we acknowledge the Collections for Human Health of the Institut Pasteur biobank (CHIP) of the Biological Resources Center of the Institut Pasteur (CRBIP) for the distribution of the blood samples. This research was funded by the Institut Pasteur, the Agence Nationale de la Recherche (ANR-21-CE15-0038-01 to P.E. and ANR-10-LABX-62-IBEID to C.B.); the Fondation pour la Recherche Médicale (FRM; EQU202503020066 to C.B.); the Programmes Transversaux de Recherche (PTR-651) from Institut Pasteur to P.E and S.A.; and the Région Île-de-France (program DIM1Health) to PBI. We acknowledge France-BioImaging infrastructure (https://ror.org/01y7vt929) supported by the French National Research Agency (ANR-24-INBS-0005 FBI BIOGEN).

## Methods

### Generation of human monocyte-derived macrophages

Blood from healthy donors was provided by the Clinical Investigation Center INVOLvE (Investigation and Volunteers for Human Health) of the Institut Pasteur. All participants received oral and written information about the research and gave written informed consent in the frame of the healthy volunteers COSIPOP cohort, after approval of the “Comité de Protection des Personnes” (CPP) Est II Ethics Committee (2023, February 20th). hMDMs isolation and differentiation were performed as previously described ^12^. Briefly, peripheral blood mononuclear cells (PBMCs) were isolated by Ficoll density-gradient centrifugation, and CD14^+^ monocytes were purified by positive magnetic selection (CD14 MicroBeads, Miltenyi Biotec). Monocytes were cultured in phenol-red-free X-VIVO 15 medium (Lonza) supplemented with 25 ng/mL recombinant human M-CSF (rh-MCSF, Miltenyi Biotec) at 37 ^*°*^C and 5% CO_2_ for 6 days. Differentiation was performed in temperature-responsive Nunc Up-Cell plates (Thermo Fisher Scientific), which allow gentle, enzyme-free detachment by transient temperature reduction, thereby preserving membrane integrity and metabolic state of the macrophages prior to seeding for infection.

### Bacterial strains and growth conditions

*Legionella pneumophila* strain Paris wild-type (WT) and its isogenic Δ*dotA* mutant, which lacks a functional Type IV secretion system (T4SS) and is unable to translocate effectors, were used in this study. Both strains carried a plasmid encoding constitutive mCherry and apramycin resistance. Bacteria were routinely cultured on ACES-buffered charcoal-yeast extract (BCYE) agar plates supplemented with 15 μg/mL apramycin at 37 ^*°*^C for 3 days. For liquid culture, colonies were inoculated into 10 mL of buffered yeast extract (BYE) medium containing apramycin (15 μg/mL) at an initial OD_600_ of 0.1 and incubated overnight at 37 ^*°*^C with shaking (200 rpm). Cultures were harvested at the late exponential phase (OD 4.2) and diluted in cell culture medium to reach the desired multiplicity of infection (MOI). Heat-killed (HK) *L. pneumophila* were freshly prepared from the same overnight WT cultures. For that, bacteria were pelleted by centrifugation (3,000 *×g*, 10 min), resuspended in sterile PBS, incubated at 70 ^*°*^C for 30 min in a water bath, and immediately transferred to ice. To avoid turbidity artefacts caused by heat-induced cellular debris, OD_600_ values used for MOI calculations were obtained from non-heat-treated aliquots. Inactivation of HK bacteria was confirmed by the absence of colony formation upon plating on BCYE agar, and structural integrity was verified by transmission electron microscopy (Extended Data Fig. 2a).

### Electron microscopy

Samples were fixed using 2.5% glutaraldehyde in 0.1 M Cacodylate buffer (pH 7.4) overnight at 4 ^*°*^C, washed for 5 min three times in 0.1 M Cacodylate buffer (pH 7.2), postfixed for 1 h in 1% osmium and rinsed with distilled water. Cells were dehydrated through a graded ethanol series followed by critical point drying with CO_2_. Dried specimens were gold/palladium sputter-coated with a gun ionic evaporator SAFEMATIC. Images were acquired using a Zeiss Crossbeam 350 field emission scanning electron microscope operating at 5 kV.

### Macrophage infection

Indium tin oxide (ITO)-coated conductive glass slides (LaserBio Labs) assembled with 8-well chamber systems (Millicell EZ, Merck Millipore) were rinsed with 100% ethanol, air-dried, and coated with 50 μg/mL poly-D-lysine (Gibco) for 1 h at room temperature, rinsed with Milli-Q water, and air-dried for 2 h. This coating procedure was independently validated to ensure homogeneous central cell distribution and confirmed not to interfere with MALDI-MSI signal acquisition. hMDMs were detached from UpCell plates and seeded at 85,000 cells per well in 500 μL of X-VIVO 15 medium and incubated overnight to reach 85% confluence. Infections were performed at MOI 10 for WT and Δ*dotA*, and an equivalent MOI 10 for HK bacteria. The MOI for HK infection was optimized by livecell imaging over 14 h, evaluating vacuolar dynamics, intra-cellular bacterial number per cell, and Annexin V positivity across initial quantities of HK bacteria to be comparable to WT-MOIs of 10, and 20. HK MOI 10 best recapitulated the number of intracellular bacteria compared to WT, while maintaining macrophage viability comparable to non-infected controls (Extended Data Fig. 2b, c). To synchronize phagocytosis, plates were centrifuged at 200*× g* for 5 min, incubated in a 37 ^*°*^C water bath for 5 min, and then transferred to a 37 ^*°*^C, 5% CO_2_ incubator for 25 min. Cells were subsequently washed three times with X-VIVO 15 for the WT and Δ*dotA* conditions, and six times for the HK. At 5 h post-infection, cells were stained with Hoechst 33342 (200 ng/mL) for 30 min at 37 ^*°*^C and immediately fixed. Hoechst staining at this concentration was validated to be compatible with MALDI-MSI spectral acquisition. Infection efficiencies of approximately 70% have been reported for this experimental model in previous work from our laboratory.

### Sample preparation for MALDI-MSI

Cells were fixed in 4% paraformaldehyde in PBS for 5 min at room temperature. Slides were then washed three times with freshly prepared 150 mM ammonium acetate, selected over PBS to preserve osmotic balance while minimizing residual salt deposition that would otherwise produce ion suppression and salt-cluster interference during MALDI ionization; this choice was empirically validated against PBS control washes. Slides were air-dried for 30 min. Tungsten pencil marks were applied to the slide edges to facilitate slide orientation across imaging platforms but were not used as references for image registration. Slides were stored at 4 ^*°*^C in a sealed, desiccated container and processed by MALDI-MSI within one week.

### MALDI mass spectrometry imaging acquisition

1,5-diaminonaphthalene hydrochloride (1,5-DAN·HCl) matrix was prepared at 4.5 mg/mL by dissolving 20 mg in 244 μL of 1 M HCl, 1,978 μL of deionized water and 2,222 μL of absolute ethanol. The solution was vortexed 30 sec and sonicated 2 min between each dilution step. Matrix was deposited uniformly using an HTX M3+ automated sprayer (HTX Imaging, Carrboro, NC, USA) with the following parameters: temperature 90 ^*°*^C; 8 passes; flow rate 50 μL/min; velocity 1,350 mm/min; track spacing 1.5 mm; CC pattern; pressure 10 psi; gas flow rate 2 L/min; drying time 30 s; nozzle height 40 mm. 1,5-DAN is an established matrix for the negative-ion MALDI analysis of low-molecular-weight metabolites and lipid species; spray-deposition conditions were adapted from Meng et al. (2020).

Matrix-Assisted Laser Desorption/Ionization – Mass Spectrometry Imaging (MALDI-MSI) was performed on a timsTOF fleX mass spectrometer (Bruker Daltonics, Bremen, Germany), with data acquired using timsControl 6.0 and FlexImaging 7.5 (Bruker Daltonics). Acquisition was performed in negative ion mode over the mass range *m/z* 50– 1,000, with laser shots set to 400 at 10 kHz. The predominant ion species were deprotonated molecules [M*−* H]^−^; chloride adducts [M+Cl]^−^ were also considered where relevant. External calibration of the TOF analyser was performed in negative-ion mode using ESI-generated sodium formate cluster ions (14 reference masses, *m/z* 112.99–996.82), yielding ~40,000 resolving power at *m/z* 500, a root-mean-square residual of 0.36 ppm and a maximum residual of 0.68 ppm (calibration-acceptance threshold 1.5 ppm); no lock-mass correction or in-batch recalibration was applied. Raw data were acquired in profile mode and subsequently exported in centroid mode (peak-picking) and converted to imzML format using ScilsLabs 2026b (Bruker Daltonics).

The laser raster step size was set to 20 μm, an empirically determined optimum balancing single-cell scale spatial resolution against ion intensity. Step sizes of 10 μm and 30 μm were also evaluated but were not retained, as 10 μm resulted in lower intensity, while 30 μm provided insufficient spatial definition. Each well was scanned over an area of approximately 4.2 mm^2^, yielding a median pixel count of ~10,500 pixels per well. Regions of interest (ROI) were guided by pre-MALDI fluorescence images and selected based on cell density, morphological integrity, and clear nuclear and bacterial signals.

### Fluorescence imaging and co-registration

Each slide was sequentially imaged in three phases to preserve technical and biological continuity from the cellular state to the final ion map: *pre-MALDI, MALDI*, and *post-MALDI*. We adopted the nomenclature used in previous approaches ^13^. Pre-MALDI imaging was performed on a CellDiscoverer 7 microscope (Carl Zeiss, Oberkochen, Germany) and combined low-magnification overviews for slide-level quality control with high-resolution fields capturing nuclear (Hoechst 33342) and bacterial (mCherry) fluorescence (see Extended Data Fig. 1). These images were used to (i) annotate each cell as infected, bystander, or non-infected; (ii) define ROIs for MALDI acquisition based on cell density, morphology (cell shape: cell area and eccentricity), and epifluorescence signal quality; and (iii) provide the spatial reference frame onto which MALDI ion images were subsequently aligned. Immediately after MALDI acquisition, post-MALDI brightfield images of the same fields were acquired on the same microscope to record the laser ablation pattern, which served as fiducial anchors for image registration (see *SpatialMetProfiler pipeline*, below). All three image layers (pre-MALDI fluorescence, MALDI ion images, and post-MALDI brightfield) were systematically retained and stored alongside their slide and well meta-data using the SpatialData framework ^14^.

### SpatialMetProfiler pipeline

We developed SpatialMetProfiler, a custom multimodal image registration and single-cell spectrum extraction pipeline implemented in Python ^15,16^ using Napari (https://napari.org/) and built on the SpatialData ^14^ framework which ensures FAIR storage for multi-modal spatial omics datasets. First, microscopy datasets were stitched and converted to the OME-Zarr file format using multiview-stitcher (https://doi.org/10.5281/zenodo.13151252). Subsequently, post-MALDI images were used as a common reference coordinate system. Pre-MALDI images were spatially aligned using rigid transforms, either automatically by means of a phase correlation-based approach, or manually by means of interactive landmark placement. Next, for each MALDI region of interest, the corresponding laser raster was interactively placed onto the ablation marks visible in the post-MALDI images by iteratively refining a non-linear transformation model, which accommodated spatial distortions introduced by matrix deposition. Average user interaction time for registration was <10 min per slide. Cells were segmented from the pre-MALDI Hoechst (nuclei) and brightfield (cells) channels using Cellpose (v3.1.1.2) ^17^ with the *cyto* model and curated by manual inspection. Each segmented cell was mapped to MALDI pixels classified *in silico* as single-cell spots with a coverage threshold above 40% of the spot area. The median per-cell pixel coverage was 12 pixels (range 8–18). Percell spectra were normalized by MALDI matrix ion intensity 1,5-DAN ([M *−*H]^−^ *m/z* 157.077). Feature identification was performed using the SCILS software (Bruker, v2026b pro). A blank subtraction-like method was performed by removing features identified in pixels having no cell inside keeping features showing intensity three times higher compared to empty pixels. The full pipeline source code, container image, parameter files, and tutorials are publicly available (see *Data and code availability*).

### Metabolite annotation

Metabolite annotation was performed through a multi-layered computational pipeline combining database matching, pathway-guided inference, adduct/isotope relationship detection, and literature-based text mining.

#### Initial database annotation

MALDI-MSI features were first annotated through METASPACE (https://metaspace2020.eu/) against the Human Metabolome Database (HMDB v5.0) and the CoreMetabolome database. The mass tolerance was set to *±*3 ppm and FDR to 5%. The ±3 ppm METASPACE window was set from the measured calibration performance rather than resolving power alone — approximately twice the 1.5 ppm acceptance threshold and about eight times the RMS residual — accommodating residual calibration drift, signal-intensity effects and pixel-to-pixel variability while remaining sufficiently selective for high-resolution annotation. For each *m/z* feature, the following adducts were considered for negative ion mode: [M *−*H]^−^, [M+Cl]^−^. This initial layer yielded 248 annotations (15.4% of detected features).

#### KEGG exact-mass matching

Unannotated features were matched against compounds present in human KEGG pathways (*Homo sapiens*, hsa) by computing the expected [M *−*H]^−^ *m/z* from KEGG exact masses and comparing against observed values within a 10 ppm tolerance window. Human pathway–compound links and exact masses were retrieved via the KEGG REST API (https://rest.kegg.jp/). Best-hit selection used a composite score balancing mass accuracy and biological context: score = (1/ppm_error) *×* log_2_(n_pathways + 1), where n_pathways is the number of KEGG pathways containing the compound. A tiered confidence system was applied: matches *≤*5 ppm was classified as higher mass-accuracy candidates, and 5–10 ppm as medium confidence. This layer contributed 49 additional annotations (3.1%).

#### Pathway-guided reverse annotation

To further annotate features, a reverse-matching approach was applied: for each KEGG compound in human metabolic pathways with a known exact mass, the expected *m/z* was calculated for five MALDI-compatible adducts — [M*−* H]^−^, [M+Cl]^−^, [M *−*H_2_O*−* H]^−^, [M+Na 2H]^−^, and [M+K*−* 2H]^−^ — and matched against remaining unannotated features within 10 ppm (tiered confidence as above). Best-hit selection among multiple candidate matches was performed using composite scoring, and annotations from enriched pathways (mummichog P < 0.15) were prioritized. This layer contributed 186 annotations (11.6%).

#### Adduct and isotope cluster detection

Relationships between detected features were explored by searching for characteristic mass differences indicative of ^13^C isotopes (+1.003), adduct conversions (e.g., [M *−*H]^−^*→* [M+Cl]^−^, Δ = 35.977 Da) and in-source neutral losses (H_2_O, CO_2_, NH_3_, H_3_PO_4_, hexose). When an unannotated feature was linked to an annotated one by an adduct conversion or neutral loss, it was annotated as an alternative ion of the same compound, extending an existing annotation rather than adding an independent metabolite. Isotope-cluster detection did not itself generate annotations; matching the expected ^13^C isotopic pattern was used only to increase confidence in these adduct-based assignments and in the underlying molecular formula. This isotope layer therefore refined and supported existing annotations, and adduct-derived alternative-ion assignments were folded into the primary annotation total rather than counted as an additional independent layer. This contributed 136 annotations (8.5%).

#### Legionella pneumophila KEGG matching

To identify potential bacterial-origin metabolites in infected macrophages, features were independently matched against KEGG compounds assigned to *L. pneumophila* (organism code: lpn) metabolic pathways, using the [M *−*H]^−^ adduct at 10 ppm tolerance with tiered confidence and composite scoring. Matches unique to the bacterial metabolome (absent from human pathways) were flagged for biological interpretation but not merged into the primary annotation table.

#### FORUM-DiseasesChem knowledge-graph annotation

To leverage published biological context, the FORUM-DiseasesChem knowledge graph (https://github.com/eMetaboHUB/Forum-DiseasesChem), an open knowledge network containing >8 billion RDF triples linking PubChem compounds, PubMed publications, and MeSH terms, was queried via its SPARQL endpoint (https://forum.semantic-metabolomics.fr/sparql). Seven MeSH terms relevant to the experimental system were used: *Legionella pneumophila* (D016952), *Legionella* (D007875), Legionnaires’ Disease (D007877), Macrophages (D008264), Macrophage Activation (D008262), Phagosomes (D010588), and Phagocytosis (D010587). Cross-graph SPARQL joins retrieved PubChem compound identifiers (CIDs) co-cited with these MeSH terms in PubMed literature. CIDs were resolved to KEGG compound identifiers via the PubChem REST API (https://pubchem.ncbi.nlm.nih.gov/rest/pug), and exact masses were obtained from KEGG or PubChem.

Remaining unannotated features were matched against these literature-derived compounds using the same five MALDI-compatible adducts and tiered ppm thresholds as above. A relevance scores weighted compounds appearing in both *Legionella* and macrophage/phagocytosis literature (3x) over pathogen-only (2x) or host-only (1x) associations. This layer contributed 34 annotations (2.1%).

#### Annotation summary and confidence

Of 1,606 *m/z* features detected across all conditions in the 50–1,000 Da range using negative ion mode acquisition, 653 (40.7%) received a putative annotation after all layers: 248 from METAS-PACE database matching (15.4%), 49 from KEGG exactmass matching (3.1%), 186 from pathway-guided reverse annotation (11.6%), 136 from adduct/isotope cluster inference (8.5%), and 34 from FORUM literature text mining (2.1%), with adduct- and isotope-cluster relationships used to support and extend these assignments rather than as an independent layer. METASPACE annotations were FDR-controlled (accurate mass, theoretical isotope-pattern agreement and spatial colocalisation of isotopic ion images), whereas KEGG- and FORUM-derived matches were accurate-mass-based only, without independent isotopic evaluation or FDR control; none resolve structural isomers sharing the same molecular formula. Annotations were assigned a confidence level following the Metabolomics Standards Initiative (MSI) reporting framework ^18^. Because all annotations rely on high-resolution accurate mass and isotope-pattern matching without complementary MS/MS confirmation, metabolite annotations reported in this study correspond to MSI Level 2 (putatively annotated compounds, based on accurate mass and isotopic pattern matched against database entries) or MSI Level 3 (putatively characterized compound classes) and should be regarded as putative pending orthogonal confirmation by compound-specific MS/MS and authentic standards. Annotations were further reviewed manually to remove implausible matches (e.g., reagent contaminants identified via FORUM such as periodate or SDS), and the curated annotation table is provided as Extended Data Table 1.

#### Untargeted LC-MS/MS metabolomics

To assess whether the metabolites putatively annotated by MSI are also detectable by an orthogonal, more established mass-spectrometry approach, untargeted LC-MS/MS metabolomics was performed on bulk extracts of hMDMs from the same four conditions (NI, WT, Δ*dotA*, HK) in a second human donor. Briefly, chromatographic separation was performed on a Thermo Fisher liquid chromatography system coupled to an Exploris 240 mass spectrometer. An Agilent HILIC column (150 *×* 2.1 mm, 2.7 μm) was maintained at 15 ^*°*^C, with a gradient elution at 0.3 mL/min using mobile phase A (H_2_O, 20 mM ammonium acetate, 5 μM medronic acid, pH 9.3) and mobile phase B (100% acetonitrile). The mass spectrometer operated in negative ion mode with a source temperature of 325 ^*°*^C, a spray voltage of 2,700 V, and sheath and auxiliary gas flows set to 55 and 10, respectively. Data was acquired using a data-dependent acquisition (DDA) method, alternating between a full MS1 scan at 120,000 resolution (at *m/z* 200) and MS2 fragmentation of the 20 most intense precursor ions.

Compounds identified against authentic standards were matched to the MSI annotation list at the level of the molecular formula; accurate-mass agreement between platforms is reported only as a concordance metric, not as the matching criterion, as the two analysers differ in mass accuracy and *m/z* range. Where a metabolite has positional or stereochemical isomers or was detected as a different adduct on the two platforms, identity was established at the level of the molecular formula and authentic standard and such cases are flagged accordingly, as isomers are not resolved by accurate mass alone. However, this LC-MS/MS analysis served solely as an orthogonal check in the presence of the annotated metabolites and was not intended for quantitative comparison between the two modalities. This important distinction is due to fundamental differences between the platforms. MSI resolves metabolites at single-cell resolution within a spatially heterogeneous population, whereas LC-MS/MS measures population-averaged bulk extracts. Given these differences in spatial resolution, ionization, and detected adducts, the LC-MS/MS data was therefore not used as a quantitative validation of the spatially resolved MSI trends. The full correspondence between MSI annotations and LC-MS/MS identifications (*m/z*, molecular formula, adduct, standard identifier and isomer/adduct caveats) is provided as Extended Data Table 2.

### Differential abundance analysis

Per-cell, 1,5-DAN-normalised spectra ([M *−*H]^−^ *m/z* 157.077) were used as input for pairwise comparisons across the four experimental conditions (NI, WT, Δ*dotA*, HK) using two complementary approaches. First, the limma package (R/Bioconductor, v3.67.3) was applied to log_2_-transformed intensities, fitting linear models with empirical Bayes moderation (eBayes); multiple-testing correction was performed using the Benjamini-Hochberg false discovery rate (FDR), and features with FDR < 0.10 were considered differentially abundant. Second, as a non-parametric, rank-based alternative robust to outliers, RankProd (R/Bioconductor, v3.37.0) was applied in parallel to the same input matrices, with significance thresholded at percentage of false positives (PFP) < 0.25. Features detected as differentially abundant by either method are reported, with priority given to those concordant between limma and RankProd. The full set of differential features per comparison is provided as Extended Data Table 3.

### Pathway representation of differentially abundant features

To relate the differentially abundant features to metabolic pathways (Extended Data Fig. 5), each feature reaching significance in a given comparison (limma, FDR < 0.10, | foldchange| > 1.5) was mapped to KEGG metabolic pathways by exact-mass matching of its *m/z* to KEGG compounds ([M *−* H]^−^ adduct, 10 ppm tolerance). For each comparison, and for each direction (the condition in which the features were elevated), we recorded the number of differentially abundant features mapping to every KEGG pathway; pathways containing at least one such feature is displayed, with dot size indicating this feature count and dot color the condition of elevation. The same membership mapping was applied to the features defining the within-condition HDB-SCAN subclusters. This analysis is a descriptive representation of the KEGG-pathway membership of the annotated differentially abundant features, not a statistical enrichment test. Because most differentially abundant features are unannotated and most pathways are represented by a single feature, no null model, permutation or over-representation statistic was applied, and no enrichment P value should be inferred from these panels.

### Dimensionality reduction and clustering

Per-cell, 1,5-DAN-normalised ([M *−*H]^−^ *m/z* 157.077) spectra were reduced in dimensionality using Uniform Manifold Approximation and Projection (UMAP) using the SCILS Unsupervised Multivariate Analysis function. UMAP parameters were set as follows: n_neighbors = 50, min_dist = 0.01, metric = Correlation Distance. For the joint WT vs NI embedding (Fig. 1), the 831 features passing a minimum-coverage filter (detected in *≥*10% of cells in either condition) were used as input. Within-condition embeddings (Fig. 2) were generated independently for each of the four conditions using the same parameters and feature selection criteria, allowing direct comparison of the number and structure of subpopulations between conditions. Unsupervised density-based clustering of the UMAP embeddings was performed with HDBSCAN (db-scan v1.2.4), with parameters min_cluster_size = 80 for the within-condition embeddings and min_cluster_size = 300 for the joint WT-NI embedding, with min_samples left at its default. These values were chosen to detect biologically meaningful subpopulations while filtering out small noise clusters.

### Four-condition shared-fit differential analysis

As an orthogonal test of whether the reduced feature abundances in WT infection depend on the choice of avirulent reference, all four conditions were modelled jointly rather than as independent pairwise comparisons. The four per-condition feature tables (NI, WT, Δ*dotA*, HK; average intensities for each of the ten biological replicates: NI and WT, n = 3; Δ*dotA* and HK, n = 2) were merged into a common feature universe by *m/z* matching within 10 ppm, yielding 1,254 features detected across all four conditions. A single linear model with a cell-means parameterisation ( ~0 + condition) was fitted across all conditions simultaneously in limma, with empirical-Bayes moderation shrinking per-feature variances toward a common prior estimated from the pooled residual degrees of freedom (df = 6). The six pairwise contrasts (WT– NI, Δ*dotA*–NI, HK–NI, WT–Δ*dotA*, WT–HK, Δ*dotA*–HK) were then extracted from this single fit, each yielding moderated t-statistics, P values and Benjamini–Hochberg-corrected FDR; features with FDR < 0.10 and | fold-change | > 1.5 were considered significant. Under this model, significant features were recovered only for the contrasts against non-infected cells (Δ*dotA*–NI, HK–NI and WT–NI); no feature reached significance in the WT–Δ*dotA*, WT–HK or Δ*dotA*–HK contrasts. For Fig. 1c, features reaching significance in the Δ*dotA*–NI or HK–NI contrasts were therefore classified by the direction of their shared-fit fold-changes in the two WT contrasts: features with log_2_ fold-change < *−*log_2_(1.5) for WT–Δ*dotA* and < 0 for WT–HK were classed as concordantly reduced in WT (n = 76), and features positive in both as concordantly increased (n = 11). This analysis provides directional concordance across two independent avirulent references, not an independent test of significance for the effector contrast.

### Statistics and reproducibility

Pipeline setup and technical validation (poly-D-lysine compatibility, HK MOI optimization, Hoechst staining compatibility, ammonium acetate washing, laser spot size optimization) were performed using hMDMs from four healthy human donors. The MSI analyses reported in this study were obtained from a single healthy human donor. Each experimental condition (NI, WT, Δ*dotA*, HK) was analyzed as independent biological replicates (separate wells, each an independent infection), three replicates for NI and WT and two for Δ*dotA* and HK, distributed across two ITO slides that were prepared and acquired in parallel. Because all wells were processed in parallel, slide membership was not treated as a batch factor. Consistent with this, per-peak coefficients of variation across biological replicates were low within each condition (median 5–16%, with few peaks exceeding 30%), and the replicate composition of the within-condition clusters was homogeneous (Extended Data Fig. 6), arguing against a dominant technical batch effect. A total of ~6,700 cells were analyzed across the WT vs NI comparison of the total >11,000 cells. Median per-cell pixel coverage was 12 pixels (range 8–18). Technical reproducibility was assessed by the median relative standard deviation (RSD) for 86 reference metabolites across independent acquisition runs, which was 11.8%. Statistical tests are specified per analysis (see *Differential abundance analysis* and *Pathway representation of differentially abundant features* above, and the corresponding figure legends). Untargeted LC-MS/MS metabolomics as an orthogonal check on MSI-annotated metabolites was performed on bulk extracts of hMDMs from the same four experimental conditions (NI, WT, Δ*dotA*, HK) in a second human donor. Unless otherwise stated, comparisons of continuous variables across more than two groups were performed using Kruskal-Wallis followed by Dunn’s post-hoc test, with Benjamini-Hochberg correction. Comparisons between two groups used the Wilcoxon rank-sum test. All p values are two-sided. Statistical analyses were performed in Python (v3.10).

### Figure preparation

Per-cell differential abundance analysis (limma, RankProd) was performed in R (v4.6.0) using the limma (v3.67.3) and RankProd (v3.37.0) Bioconductor packages. UMAP embeddings and feature-table exports were computed in SCiLS Lab. All quantitative plots, UMAP and clustering visualizations, and heatmaps of differentially abundant features and cluster signatures were generated in Python (v3.10) using Matplotlib (v3.5.1) and Seaborn (v0.12.2). Heatmaps used seaborn.heatmap with diverging (RdBu_r) or sequential (YlOrRd) colour maps as appropriate. Statistical comparisons used SciPy (v1.10.1). Prism 11 (GraphPad) was also used to plot several graphs. Elements in Fig. 1a were created in BioRender (https://biorender.com/). MARTINEZ OCA, P. (2026) https://BioRender.com/kp98f2d.

### Generative AI statement

During the preparation of this work, authors used ChatGPT 5.5 (OpenAI), Gemini 3.1 pro (Google) and Claude Opus 4.8 (Anthropic) Large Language Models (LLMs) to correct grammar and flow of some parts of the text. Claude Code (Opus 4.6, Anthropic) via plug-in for VS Code (Microsoft) was used to generate some of the code and analysis scripts. After using these tools, authors reviewed and edited the content as needed and take full responsibility for the content of the publication.

### Data and code availability

Raw MALDI-MSI datasets, pre-MALDI fluorescence images, and post-MALDI brightfield images were uploaded to METASPACE (https://metaspace2020.eu/) and are publicly accessible under project ID ‘MetaBact-MSI’ (https://metaspace2020.org/project/MetaBact-MSI). The SpatialMetProfiler pipeline, including source code, container images, parameter files, and tutorials, is openly available at https://github.com/pescoll/MSI-Macrophages-Legionella.git under an open-source license. Analysis code (R and Python) generating the figures of this manuscript is available in the same repository. Any additional information required to reanalyze the data is available from the corresponding authors upon request.

## Extended Data

### Extended Data Figures

**Extended Data Fig. 1.**
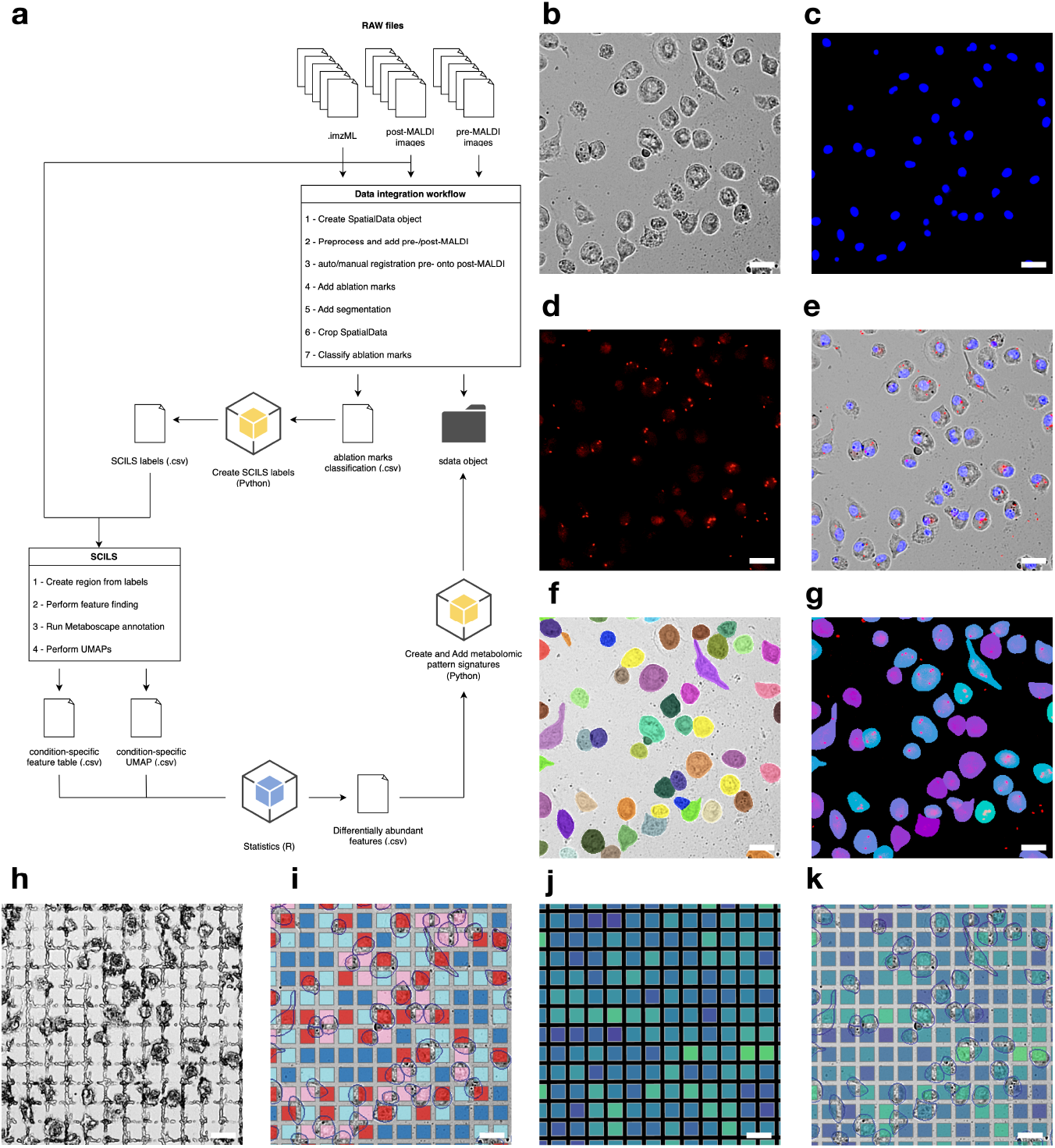
Detailed SpatialMetProfiler pipeline. Representative WT-infected field illustrating the sequential imaging, segmentation, registration and single-cell spectrum-extraction steps of the SpatialMetProfiler workflow (Methods). (**a**) Overview of the pipeline: raw MALDI spectra (.imzML) with pre- and post-MALDI images are integrated into a SpatialData object (registration, ablation-mark addition, segmentation, cropping and ablation-mark classification), single-cell spectra are extracted in SCILS, and differentialabundance and metabolic-pattern analyses are performed in Python and R. (**b**) Brightfield (BF) image of adherent hMDMs on the ITO-coated slide. (**c**) Hoechst nuclear staining of the same field, used for nucleus detection and cell counting. (**d**) mCherry fluorescence identifying intracellular *Legionella pneumophila* (mCherry-expressing), puncta correspond to bacteria. (**e**) Overlay of BF, Hoechst and mCherry channels. (**f**) Single-cell segmentation from automated image analysis (each mask, one cell). (**g**) Segmentation masks overlaid on the bacterial fluorescence, color-coded according to mCherry fluorescence (SD/mean). (**h**) Post-MALDI BF image showing the regular grid of laser ablation marks. (**i**) Ablation marks color-coded by classification: no cell (dark blue); <40% of the ablation mark covered by a cell (blue); multiple cells (pink); exactly one cell covering >40% of the ablation mark (dark red). Only single-cell marks (dark red) are retained in the downstream analyses. (**j**) MALDI ion-intensity map of the same field (L-glutamic acid). (**k**) The same ion-intensity map with single-cell segmentation masks overlaid, illustrating assignment of per-pixel intensities to individual cells. Scale bar = 25 μm.

**Extended Data Fig. 2.**
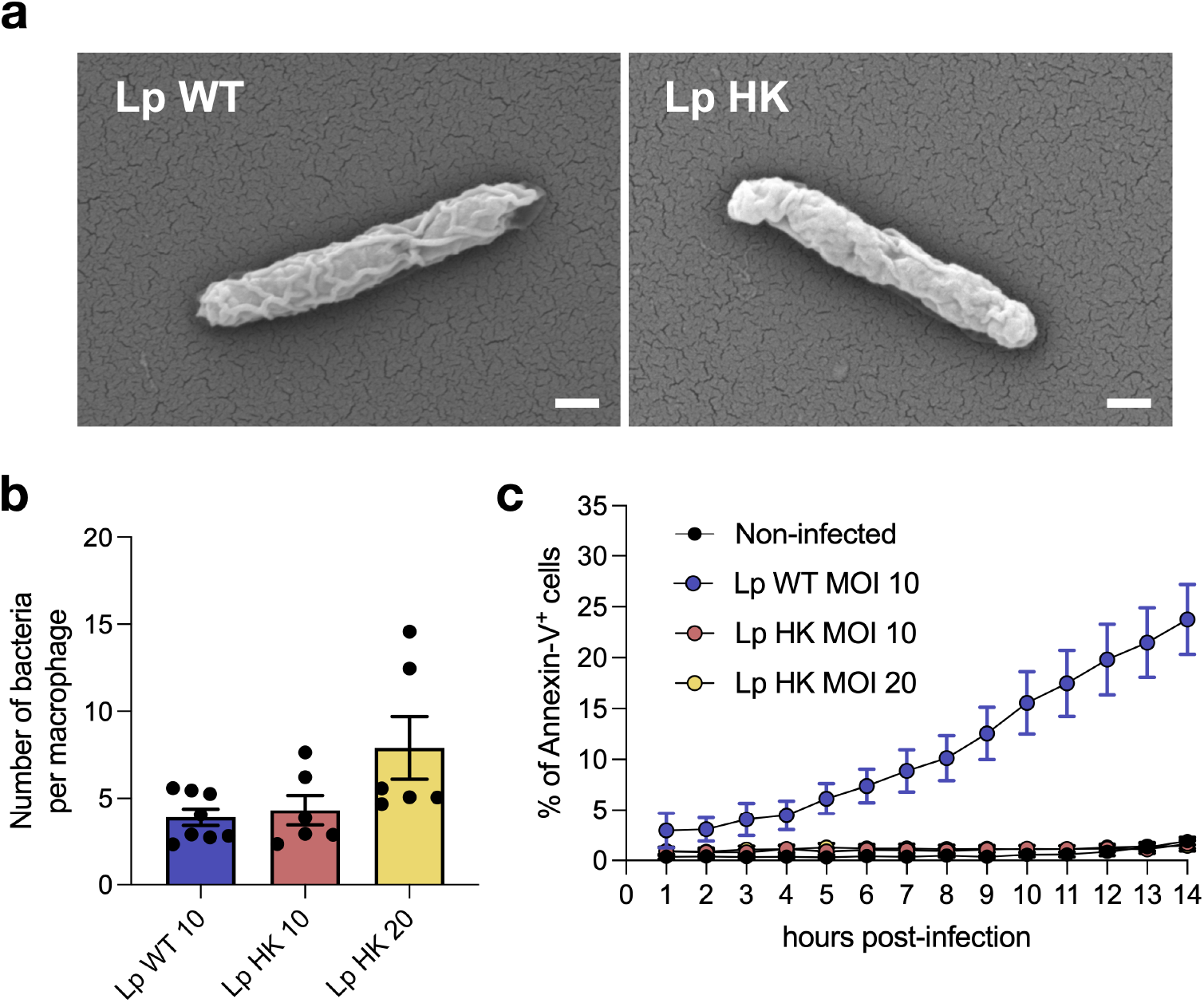
Characterization of heat-killed (HK) *L. pneumophila*. (**a**) Electron micrographs of Lp WT and Lp HK, showing that heat inactivation preserves bacterial morphology; loss of viability was confirmed by absence of colony formation on BCYE agar. Scale bar = 200 nm. (**b**) Intracellular bacteria per macrophage for Lp WT (MOI 10), Lp HK (MOI 10) and Lp HK (MOI 20) by live-cell imaging; HK at MOI 10 matches the WT MOI 10 load, whereas MOI 20 exceeds it (mean ± SEM). (**c**) Percentage of Annexin-V+ macrophages over 14 h for non-infected cells, Lp WT (MOI 10) and Lp HK (MOI 10 and 20); WT reaches ~24% by 14 h whereas both HK conditions remain at the non-infected baseline (~1–2%), confirming HK bacteria do not compromise viability (mean ± SEM).

**Extended Data Fig. 3.**
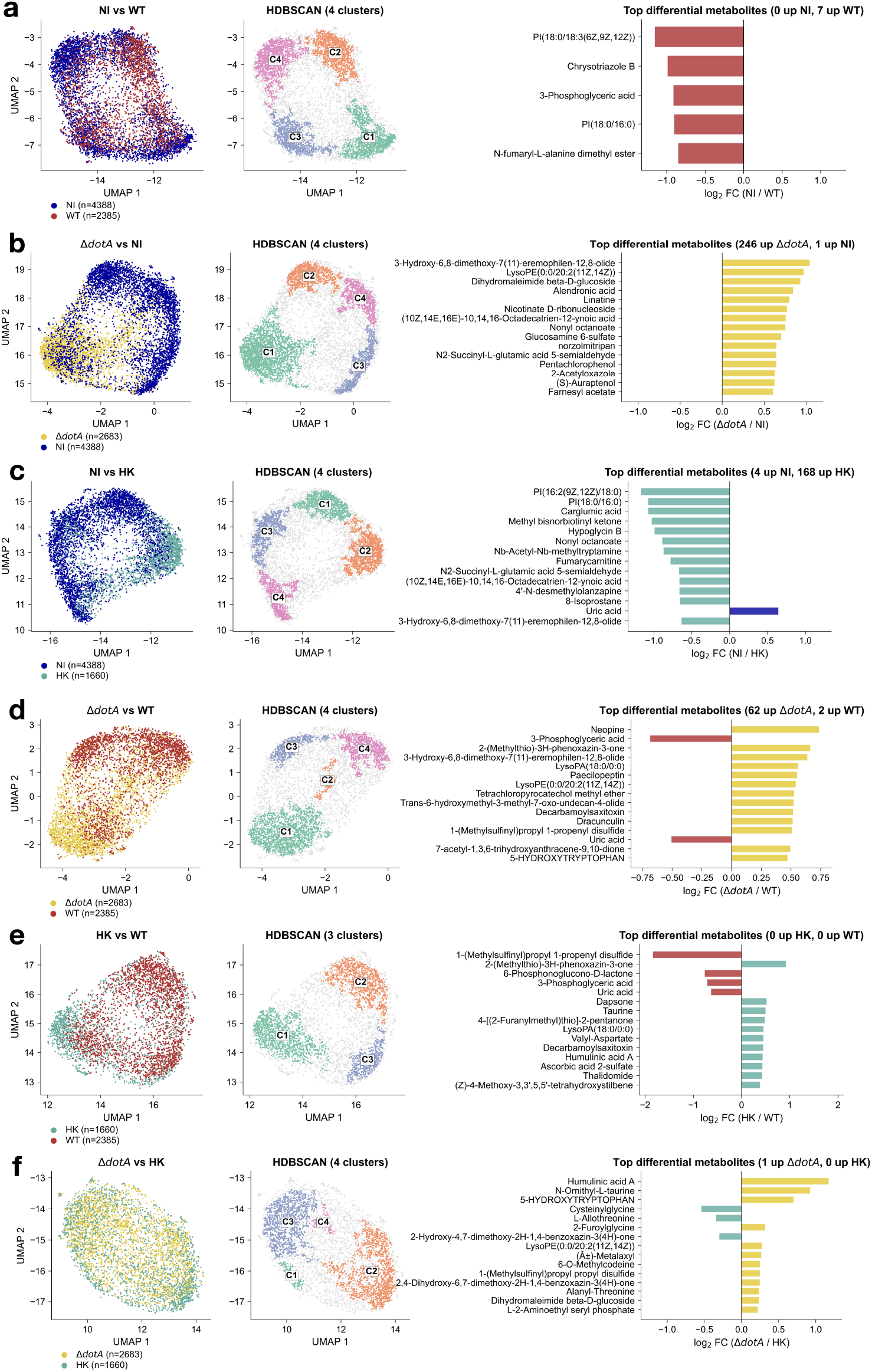
Pairwise single-cell metabolic comparisons across all six condition pairs. For each pair, a joint UMAP colored by condition (left), unsupervised HDBSCAN clustering of that embedding (middle), and the top differentially abundant features ranked by log_2_ fold-change (right, limma, FDR < 0.10), with fold-change in the direction stated. (**a**) NI vs WT: 7 features elevated in WT, none in NI. (**b**) Δ*dotA* vs NI: 246 elevated in Δ*dotA*, 1 in NI. (**c**) NI vs HK: 168 elevated in HK, 4 in NI. (**d**) Δ*dotA* vs WT: 62 elevated in Δ*dotA* (equivalently, suppressed in WT), 2 elevated in WT (the effector contrast summarized in Fig. 1c). (**e**) HK vs WT: no feature reaches significance. (**f**) Δ*dotA* vs HK: a single differential feature (5-hydroxytryptophan, slightly higher in Δ*dotA*), indicating that bacterial viability contributes little beyond uptake of dead bacteria. Panels d and e underlie the WT-suppression concordance (Fig. 1c), while panels b and c underlie the sensing/uptake concordance (Fig. 2e).

**Extended Data Fig. 4.**
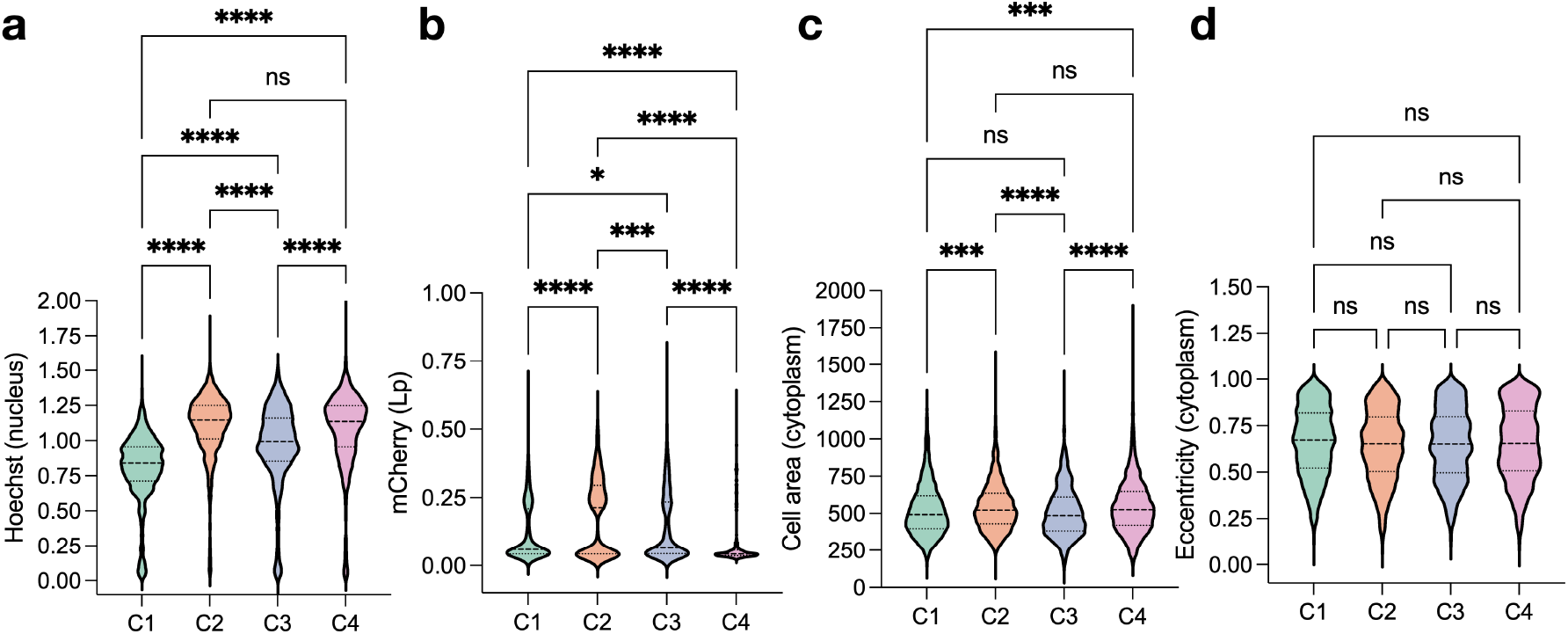
Per-cluster distributions of cellular parameters from HDBSCAN of WT-NI UMAP. (**a**) Hoechst fluorescence. (**b**) mCherry fluorescence (Lp = *L. pneumophila*). (**c**) Cell area. (**d**) Cellular eccentricity.

**Extended Data Fig. 5.**
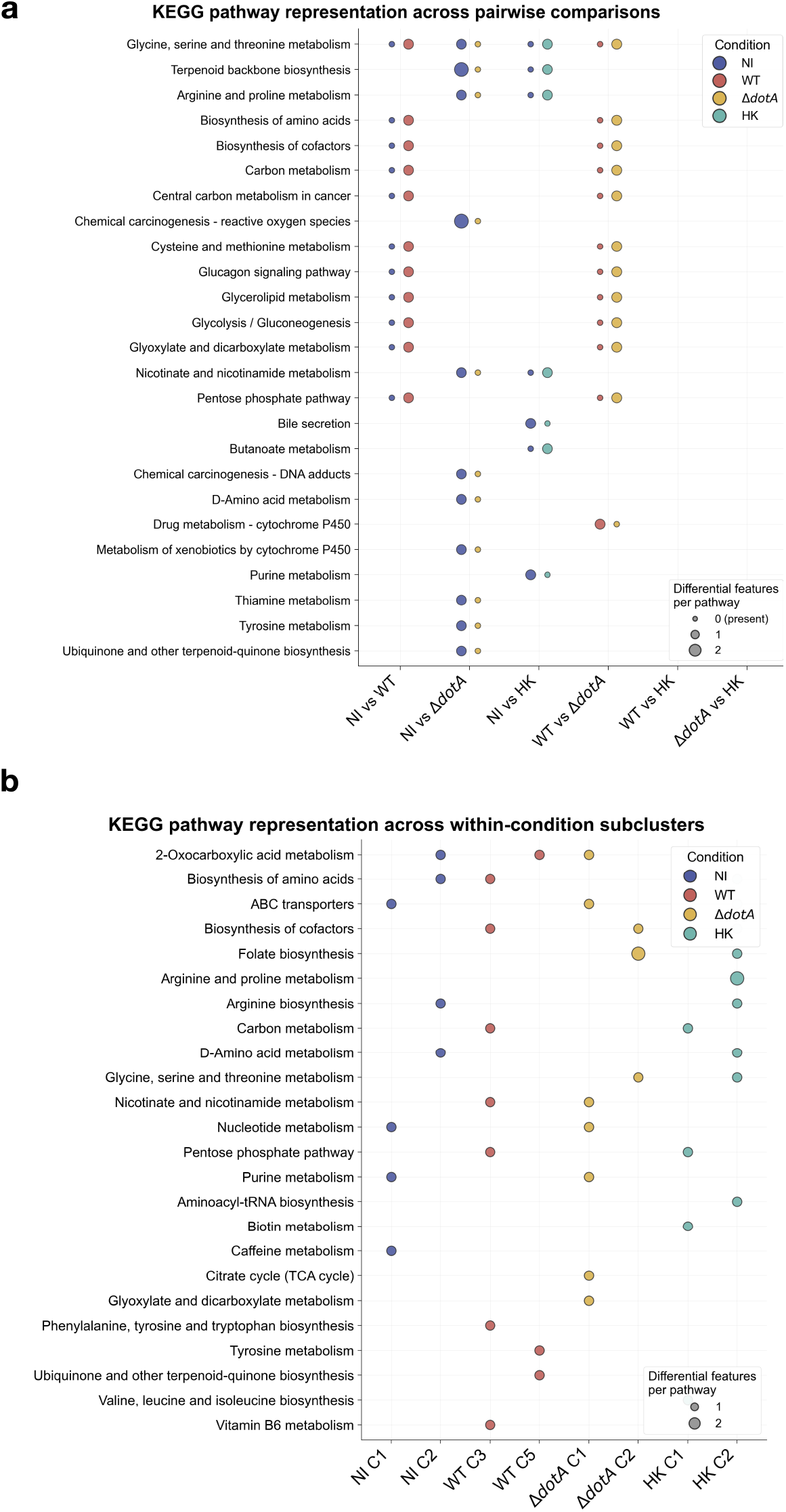
KEGG pathway representation of differentially abundant features, across pairwise comparisons and within-condition subclusters. Differentially abundant features (limma, FDR < 0.10, |fold-change| > 1.5) were mapped to KEGG metabolic pathways by exact-mass matching of their *m/z* to KEGG compounds ([M*−*H]^−^ adduct, 10 ppm tolerance; Methods). Dots mark pathways containing at least one such feature; dot colour indicates the condition in which the feature(s) are elevated (NI, blue; WT, red; Δ*dotA*, yellow; HK, green) and dot size the number of differentially abundant features mapping to that pathway. (**a**) Pathway membership for each of the six pairwise comparisons (NI vs WT, NI vs Δ*dotA*, NI vs HK, WT vs Δ*dotA*, WT vs HK, Δ*dotA* vs HK). (**b**) Pathway membership across the within-condition HDBSCAN subclusters for which annotated differential features were available (NI C1-C2, WT C3 and C5, Δ*dotA* C1-C2, HK C1-C2). This is a descriptive representation of KEGG-pathway membership, not a statistical enrichment test. No null model, permutation or over-representation statistic was applied and no enrichment P value should be inferred. Because most differentially abundant features are unannotated, only a small minority of features contribute to these panels, and almost all pathways shown are represented by a single feature.

**Extended Data Fig. 6.**
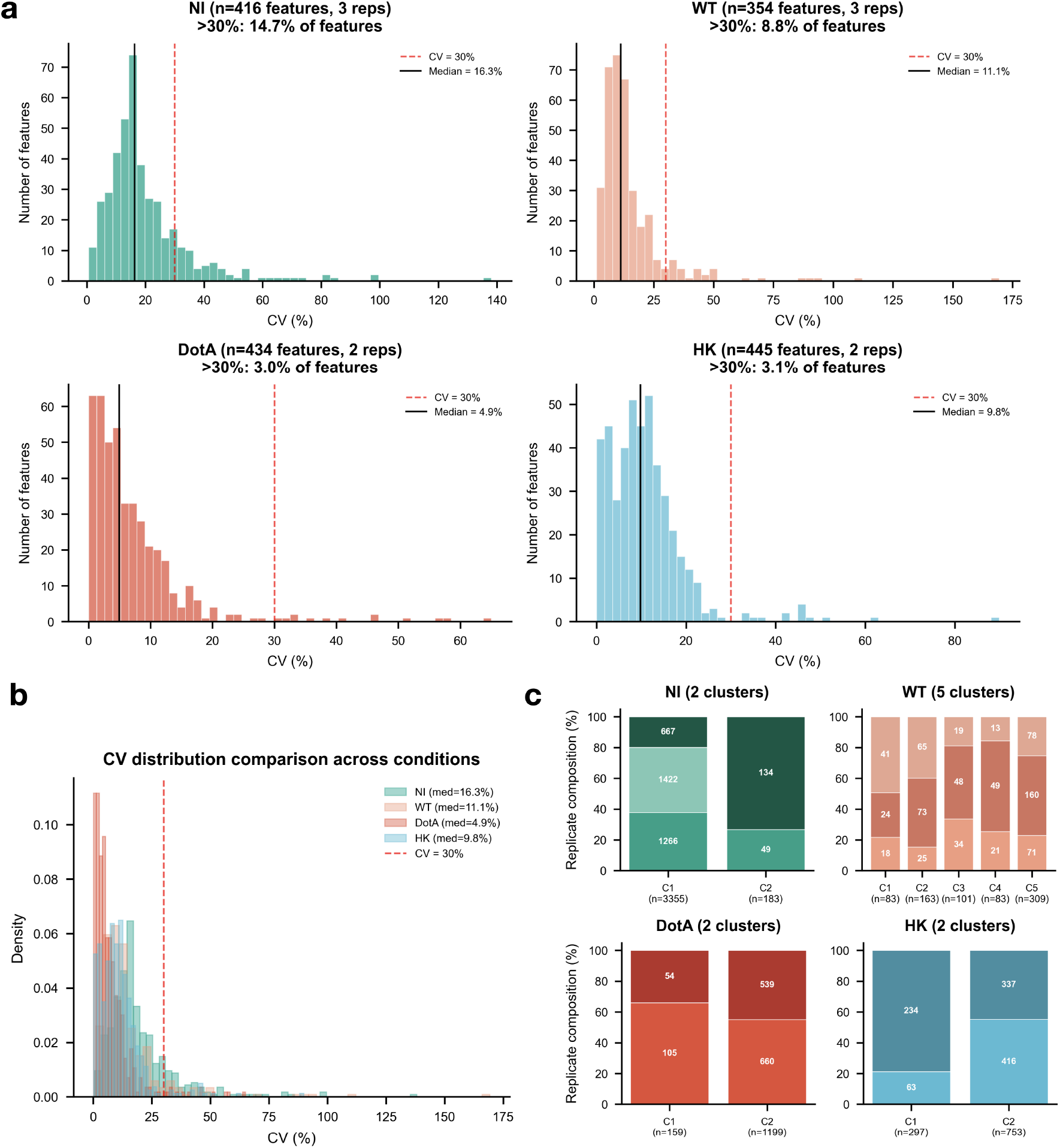
Technical reproducibility and replicate composition of within-condition subclusters. (**a**) Per-condition CV distributions of feature intensities across biological replicates (NI, WT: n = 3; Δ*dotA*, HK: n = 2); median CV and fraction with CV > 30% annotated. (**b**) Overlaid CV distributions. (**c**) Replicate composition of each subcluster, indicating low technical variability.

### Extended Data Tables

**Extended Data Table 1. Curated metabolite annotation table.** Putative annotations for the 653 annotated *m/z* features (of 1,606 detected, 50–1,000 Da, negative ion mode), listing observed *m/z*, matched KEGG/HMDB compound, adduct, ppm error, annotation layer (METASPACE, KEGG exact-mass, pathway-guided reverse, adduct/isotope inference, FORUM literature) and MSI confidence level (Level 2/3, or Level 1 where confirmed by LC-MS/MS). Provided as a separate file.

**Extended Data Table 2. Correspondence between MSI annotations and LC-MS/MS identifications.** The 33 MSI annotations independently matched to a metabolite detected and identified by untargeted LC-MS/MS against authentic standards, listing *m/z*, molecular formula, adduct, standard identifier, metabolic class, and isomer/adduct caveats. Provided as a separate file.

**Extended Data Table 3. Full set of differentially abundant features per comparison.** Differential-abundance results for each of the six pairwise comparisons (limma FDR and RankProd PFP, log_2_ fold-change, annotation, and concordance flag between methods). Provided as a separate file.

